# Structural and Conformational Impact of Deleterious Spike Protein Mutations in SARS-CoV-2 Omicron Lineages

**DOI:** 10.1101/2025.02.25.639252

**Authors:** Aqsa Khalid, Kumail Ahmed, Akbar Kanji, Tania Munir, Sayed Ali Raza Shah Bukhari, Zahra Hasan, Wesley C. Van Voorhis, Najeeha Talat Iqbal

## Abstract

**Background:** A large number of mutations in the Spike (S) protein of the SARS-CoV-2 omicron variant have been noted to alter the receptor binding domain (RBD) and increase the binding surface and enhance the opening of the binding pocket. The cumulative effect of S1 and S2 subunit mutations can influence the conformational dynamics of the binding surface, facilitating the release of viral genome into host cells.

**Aim:** This study investigates the deleterious mutations across all Omicron lineages identified in our analysis and their effect on the conformational stability of RBD opening.

**Methods:** Whole Genome Sequencing of 231 SARS-CoV-2 positive patients in Karachi, Pakistan, were performed using Illumina Miseq instrument and raw reads were analyzed using viralrecon pipeline. The mutational effects of omicron variant on the stability of S protein, including wild-type (7FG7), close (6VXX) and open (6VYB) states, were assessed through MD simulations.

**Results:** Four deleterious missense mutations (Tyr505His, Asn764Lys, Asp950Asn, Asn969Lys) were identified in the S1 and S2 subunit of the S protein of omicron variant. In the wildtype and open state mutant models, Tyr505His, Asp950Asn and Asn969Lys caused destabilizing effects, higher RMSDs vs. wild-type, and fluctuations in the RBD (438-510) region and S2 subunit (946-1010), compared to the native structure. These mutations increased the binding pocket propensity to open in mutant model compared to the native open conformation (6VYB). This structural change promoted trimer opening in the open state through α-helix movement in the S2 subunit away from the RBD region. In the closed state, only S2 subunit mutations (Asp950Asn and Asn969Lys) lead to predicted destabilization through the movement of protomer C towards protomer B (RBD region). These S2 subunit mutations are predicted to stabilize the RBD “down” conformation potentially enhancing spike antigenic heterogeneity.

**Conclusion:** This study highlighted the cumulative effect of S1 (Tyr505His) and S2 (Asp950Asn and Asn969Lys) subunits mutations on different S protein states, potentially controlling its conformational dynamics and presentation to host receptors. Future experimental studies are needed to elucidate the biological significance of these alterations, particularly by establishing a link between the identified mutations and their impact on viral biology.

## Introduction

The COVID-19 pandemic has witnessed a persistent rise in new evolving mutations, causing significant challenges and difficulties for initiatives to improve global health [1]. The evolving mutations within the Spike protein (S) of SARS-CoV-2 can potentially affect transmissibility, virulence, and evasion of the immune system. Similar to other viruses, SARS-CoV-2 also naturally undergoes mutations over time due to its highly variable mutation rate, and most of these mutations do not directly benefit the virus [2], [3], [4]. The mutations in the SARS-CoV-2 are either neutral or mildly deleterious, however, the highly deleterious mutations are less likely to propagate successfully due to evolutionary selection. Mutations that appear and are predicted to lead to mild and highly deleterious functions need further investigation for selection pressure [5]. Experimental studies such as site-directed mutagenesis through cell transfection [6], Biolayer Interferometry (BLI) S protein-ACE2 binding assay [7], and protein characterization assays [8] have primarily focused on mutations occurring in the receptor-binding domain (RBD). However, genetic alterations in other regions of the SARS-CoV-2 genome also need attention to incorporate factors not considered in cell lines transfection and S protein-ACE2 binding assays. Genetic alterations in the genome of SARS-CoV-2 present a considerable challenge to pandemic prevention and control efforts, potentially hindering progress in therapeutics, diagnostics, and vaccination [9]. Variants of Concern (VOC) enable the virus to evade immunity through various mechanisms, such as altered interactions with immune regulatory genes, epitope loss, evasion of T-cell killing, and reduced affinity for neutralizing antibodies [10], [11]. Furthermore, circulating VOCs exhibit enhanced transmissibility, infectivity, and immune evasion due to structural rearrangements that increase the accessibility and binding affinity of the receptor-binding domain (RBD) to ACE2 [12]. The differences in the interaction patterns of key amino acid residues and modifications in the binding pocket led to the larger binding interface and higher affinity to ACE2 in the RBD of SARS-CoV-2, highlighting the impact of virus evolution on immunity and vaccinations [13]. Previous studies also proposed that SARS-CoV-2 S protein has lower stability in comparison to SARS-CoV S protein, with the S2 fusion being more conserved than the S1 domain [14]. Additionally, the conformational transition of the wild-type SARS-CoV-2 S protein between its open and closed states plays a crucial role in ACE2 receptor binding. This transition occurs through the transient movement of the RBD, which alternately hides or exposes the receptor-binding determinants. These conformational changes facilitate viral fusion with the host membrane, aiding in the release of the viral genome into the host cell [15].

In this study, we aim to elucidate the impact of mildly deleterious mutations within the RBD and S2 subunit of the S protein, with a particular focus on their structural and functional consequences in the wild, closed, and open states. We specifically examine the opening of the RBD upon these mutations. The dynamics of the identified mutations in the S2 subunit have not been extensively studied using either experimental or computational approaches. Our research seeks to investigate the effects of these mutations, particularly in the S2 subunit, identified from whole genome sequencing (WGS) data of patients from Karachi, Pakistan, and how do they facilitate viral entry into host cells.

## Methodology

### Study Design

The current study is part of the United World for Antiviral Research Network (UWARN) for surveillance of SARS-CoV-2 in patients presenting clinical symptoms. The research protocol was approved by the Ethical Review Committee of Aga Khan University (ERC protocol 2021-4794) Pakistan. Patients were recruited from inpatient and outpatient settings from The Aga Khan University Hospital (AKUH) including age groups 1 to 75 years from 2021 to 2022, after informed consent. Patients were enrolled as inpatients if they were hospitalized *≤*1 week with confirmed diagnosis of the COVID-19, with or without clinical signs and symptoms consistent with the COVID-19 infection. Those who were referred by study physicians having clinical signs and symptoms consistent with the COVID-19 infection or had a history of exposure with COVID-19 patient or confirmed diagnosis using PCR or rapid Antigen test were enrolled as outpatients. The consent was taken from the enrolled participants either in-person or via telephone by the patient themselves or from their Legally Authorized Representatives. Those who were out of age criteria or unable to provide consent and had disabilities were excluded from the study.

### Sample Collection

Nasopharyngeal swab samples tested positive for SARS-CoV-2 through reverse transcription (RT) polymerase chain reaction (PCR) using the Roche Cobas 6800 assay at the Section of Molecular Pathology, AKUH, Karachi, Pakistan. The only specimen that exhibited amplification at threshold cycle (Ct) value *≤* 30 was selected for the comprehensive whole genome sequencing (WGS).

### RNA Extraction

RNA extraction of all positive SARS-CoV-2 nasopharyngeal swabs specimen was carried out using QIAamp Viral RNA Mini Kit (Qiagen, Heiden, Germany, cat: 52906) according to the manufacturer’s recommendations. RNA was eluted in 80µl of elution buffer. RNA purity and quantity were checked by using Qubit 4 fluorometer (Life Technologies, Singapore) and stored at −80°C for sequencing.

### Complementary DNA (cDNA) synthesis and Tiled PCR

cDNA was synthesized using the Superscript IV VILO™ master mix (Thermo Scientific, MA, USA) following the guidelines provided by the manufacturer [16]. The protocol was run on the Eppendorf Master Cycler X50a (Hamburg, Germany) platform. Moreover, Tiled PCR was performed using Q5 High Fidelity PCR master mix (New England Bio Lab, UK). Briefly, cDNA was used as a template and two separate PCR were set up and mixed with Q5 High Fidelity master mix along with specifically designed Primer Pool A and Pool B by ARTIC nCoV 2019 network_V3 by IDT (USA)[17] with a concentration of 10µM of each, creating a total reaction volume of 25µl respectively. Cycling conditions includes denaturation at 98 °C for 30 sec, followed by 35 cycles at 95 °C for 15 sec, and 63 °C for 5 min, and finally hold at 4 °C. The PCR protocol was executed on the Eppendorf Master Cycler X50a (Hamburg, Germany) platform. Two PCR amplicons were generated; equal volumes of both were combined, and the total DNA concentration was determined using the Qubit HS DNA assay (Invitrogen).

### Library Preparation

Amplicon Library was generated using the Illumina DNA Library Prep Kit (Illumina, Portland, USA) according to manufacturer’s instructions [18], using 1 ng of input DNA. Tagged PCR samples were purified with 30 µL of AMPure XP beads (Beckmann Coulter, CA, USA). Sample library normalization and MiSeq sample loading were carried out according to the Illumina Library Prep protocol. All normalized libraries were pooled and spiked with Phi-X control at a final concentration of 0.1 pM followed by loading onto a Mi-Seq reagent kit V3 600 cycle (Illumina, Portland, USA) [19] and sequenced on a MiSeq Platform (Illumina, Portland, USA).

### Sequencing Analysis

Raw reads from Illumina sequencing were assembled using Viralrecon [20]and results were further validated using in-house pipeline. The protocol includes the quality check of raw reads using FASTQC [21] and removal of low-quality reads through trimmomatic [22]. Reads are then sorted against the reference SARS-CoV-2 genome (NC_045512.2) using SAMtools [23]. Variants were called and their allele frequency were calculated using Vcftools [24] and genetic annotations were identified using SNPEff [25]. The SNPEff tool classify the mutations to be high (frameshifts), moderate (missense variants, in-frame insertions or deletions) and low (synonymous variants or in introns or UTRs) impact mutations. We considered moderate-impact mutations identified to further analyze their potential effects. The SIFT software [26] was employed to classify moderate-impact mutations as deleterious (SIFT score *≤* 0.05) or neutral/non-deleterious (SIFT score > 0.05). Mutations predicted to be deleterious were subsequently selected for downstream analysis. In addition, lineages were also assigned based on established dynamic lineage classification method proposed by Rambaut et al. via the Phylogenetic Assignment of named Global Outbreak LINeages (PANGOLIN) software suite (https://github.com/hCoV-2019/pangolin) and deposited data into GISAID (https://www.gisaid.org/).

### Homology Modeling

Cryo-EM three-dimensional (3D) wild-type structure (PDB code: 7FG7) [27], open (PDB: 6VYB) and closed (PDB: 6VXX)[28] conformations of SARS-CoV-2 spike glycoprotein (S) were retrieved from RCSB Protein Data Bank. The identified mutations from WGS in the S1 and S2 subunit of spike protein were modeled using Modeller v.10.3 [29]. To perform the homology modeling Multiple Sequence Alignment (MSA) of the targets (ID: P0DTC2) and template sequences were performed using T-Coffee [30]. A total of 20 homology models of S protein per subunit were generated and assessed on the basis of ERRAT[31] and PROCHECK scores[32]. ERRAT shows the overall quality of the predicted 3D structure and PROCHECK asses the stereochemistry properties by phi (ϕ) - psi (ψ) torsion angles through Ramachandran plot. Furthermore, in order to obtain the most stable structural conformation, energy minimization was performed using Nosé-Poincaré Andersen (NPA) dynamics algorithm of MOE at a time step of 0.002 for 500 ps[33]. After the energy minimization, MD simulation of the final selected mutant model was performed through using GROMACS.

### Molecular Dynamic (MD) Simulations

MD simulations of energy minimized mutant S protein models were performed for 100 ns using Gromacs software version 2018 [34]. The mutated protein structures were solvated with TIP3P water molecules in a cubic box, and counterions were added to neutralize the system. The systems were then energy minimized using the steepest descent and conjugate gradient algorithms to remove steric clashes and close contacts. The equilibration of the systems was carried out in the NPT ensemble for 100 ps, during which the temperature and pressure were maintained at 300 K and 1 bar, respectively. The production MD simulations were carried out for 100 ns using the CHARMM36 force field[35]. The trajectories generated from the MD simulations were analyzed to obtain Root-Mean Square deviations (RMSD) of protein-ligand complexes to investigate their conformational stability.

## Results

The current study is part of the UWARN project aimed at performing whole-genome sequencing of SARS-CoV-2 to explore genetic variations. A total of 231 SARS-CoV-2 samples, collected from various districts of Karachi between May 2021 and October 2022, were sequenced using NGS (illumina, MiSeq platform) at Aga Khan University, Pakistan. Out of the 231 sequenced samples, 165 isolates (Suppl. Appendix I, S2) were deposited into the GISAID database. The variant distribution across these isolates revealed a predominance of omicron variants (106; 64%), followed by Delta (31; 19%), Beta (3; 1.8%) and Alpha (4; 2.5%). Due to low sequence coverage, the lineages for 21 isolates (12.7%) could not be assigned. The omicron lineage distribution (n=106) revealed BA.5 [63 (59.4%)] as the most abundant strain, followed by BA.2 [18 (17%)], BA.1 [14 (13.2%)], BA.3 [4 (3.8%)], BA.4 [3 (2.8%)], BV.2 [3 (2.8%)], and CK.1.3 [1 (0.94%)] as shown in Figure 1. Omicron isolates having coverage (>90%) and sufficient depth (>10x) was considered to explore the mutations in the S1 and s2 subunit of Spike (S) protein (Suppl. appendix I, S2).

**Figure 1:**
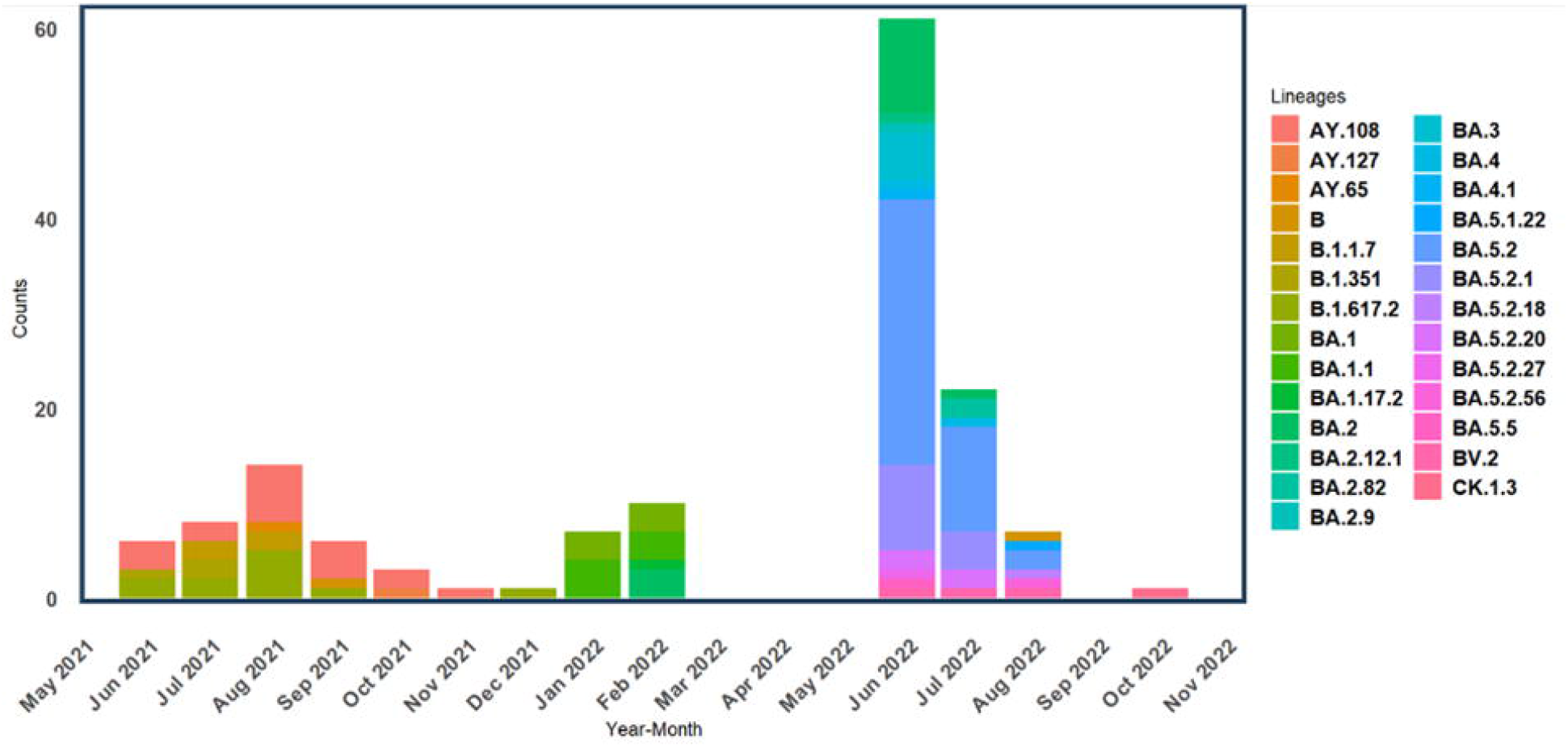
Distribution of Pangolin lineages of SARS-CoV-2 variants. The predominant variants observed are within the omicron lineage, specifically BA.1,BA.2,and BA.5.

### Missense Mutations and associated patient profiles in the Sequenced SARS-CoV-2 Genomes

Our sequencing results revealed multiple single nucleotide polymorphisms (SNPs) in the ORF1a, ORF1b, spike glycoprotein (S), ORF3a, ORF6, ORF7a, ORF8, ORF10, envelope protein (E), membrane protein (M) and nucleocapsid protein (N) regions. These mutations play an important role in adapting SARS-CoV-2 to the human host. Few mutations were identified in the E, M, and ORF genes, and were supposed to be non-deleterious and did not significantly affect protein structure due to their SIFT score > 0.05. The impact of mutations on SARS-CoV-2 proteins, including S, E, M, and N, was assessed using SIFFT software. However, mutations affecting the conformation of the Spike glycoprotein (S) had significance in terms of viral pathogenesis [36].

Notably, omicron isolates (n= 52) had 10 missense mutations with deleterious impact observed within the S protein, including Lys212Ser in the N-terminal domain (NTD), Tyr505His in the Receptor Binding Domain (RBD), Tyr716Leu and Tyr741His, Asn764Lys in the S1/S2 cleavage site, and Ser939Phe and Asp950Asn in the Heptapeptide repeat sequence 1 (HR1), as well as Asn969Lys, Glu1151Asp, and Val1128Ala between HR1 and HR2 of the S2 subunit (Figure 2). Four mutations (Tyr505His, Asn764Lys, Asp950Asn, Asn969Lys) with variant allele frequency (VAF) of >0.9 and higher prevalence in our samples. Tyr505His was identified in 30/52 samples with a VAF of 1.0, Asn764Lys in 47/52 samples with a VAF of 0.99, Asp950Asn in 3/52 samples with a VAF of 0.95, and Asn969Lys in 46/52 samples with a VAF of 0.99 (suppl. appendix sheet 3, Figure S1). We initially excluded mutations with a variant allele frequency (VAF) of <0.9, such as Tyr741Ser, Glu1151Asp, and Val1128Ala (Figure S1). Subsequently, we removed low-prevalence mutations, including Leu212Ser and Ser939Phe, found in one sample, and Thr716Leu, identified in two samples.

**Figure 2:**
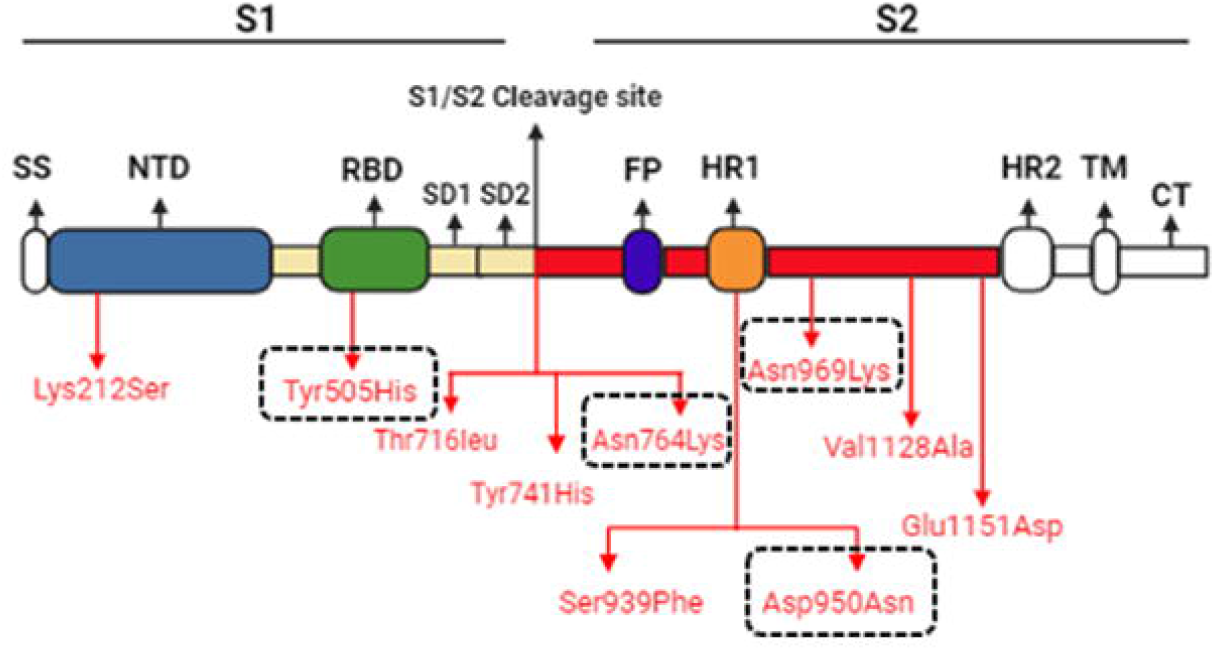
Deleterious mutations identified in the S1, Furin region, and S2 subunit of the spike glycoprotein (S). The black dashed box represents mutations with VAF (>0.9) and higher prevalence in our samples

The homology model of these mutations was built for wild-type, open and closed conformations of spike-glycoprotein to analyze the conformational stability and whether these mutations in the open and closed state favor accessibility for binding of ACE2 receptor and viral entry into the host cell.

The demographic and clinical profiles of these 52 selected omicrons having missense deleterious mutation cases are summarized in Table 1. Approximately half of the patients were female (48%) with mean age of 38 years for both genders (male and female) and all of them belongs to non-severe cases category. However, these cases had other associated symptoms including fever (52%), cough (25%), shortnes of breath (5.8%), sore throat (9.6%) and body aches (7.7%). At the time of enrollment, 43(82.6%) received both 1^st^ and 2^nd^ doses of vaccination and majority of the cases received Sinopharm vaccine (Table 1).

**Table 1:**
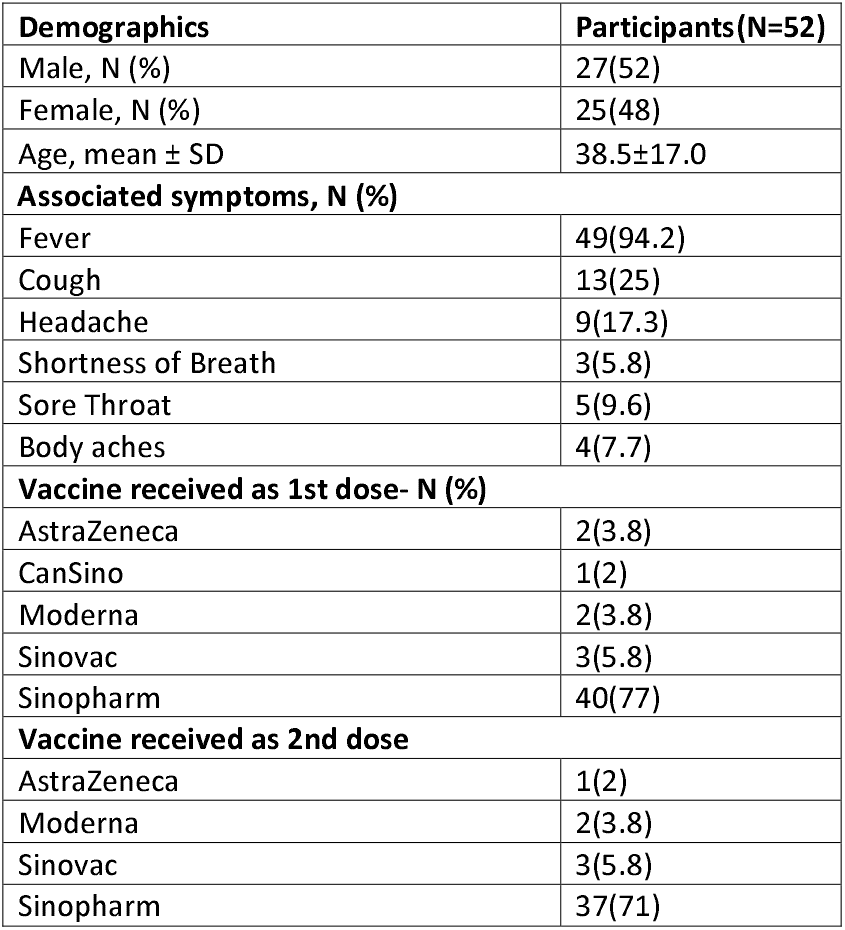
Clinical characteristics of patients with Omicron variant COVID-19 in Karachi, Pakistan.

### Structural and Functional Modeling of SARS-CoV-2 Mutations

The sequencing analysis revealed 08 deleterious mutations in the S protein as discussed earlier (Figure 2). The His505 lies in the RBD region of the S1 subunit of the protein. Other mutations in circulating strains were found in the Furin cleavage and S2 subunit. To assess the effect of these mutations, we created the mutants His505, Lys764, Asn950, and Lys969 for the wild, closed, and open conformations of S protein. The mutant models of the wild-type (7FG7) closed (6VXX) and open (6VYB) crystallographic structures selected for downstream analysis showed ERRAT scores of 55.5, 61.47, and 65.59, respectively. The Ramachandran plot (Figure S2) indicates a minimal number of residues for 7FG7 (1.5%), 6VXX (0.5%), and 6VYB (0.2) in the disallowed region (Table S1). ERRAT scores (>50) and minimal residues (<2%) in disallowed regions in all the mutant models indicate that these models are reliable and could be used for downstream analyses. In addition, two sets of MD simulations were performed at 100 ns for each mutant and native model of wild-type, closed, and open conformations. The most stabilized structure from the last energy frame was extracted to compare the dynamics of the structural conformations, root means square deviation (RMSD) and root mean square fluctuations (RMSF) profile.

Starting from the wild-type protein, the observed mutations are predicted to impose structural changes, with the superposition of native and mutant structures showing an RMSD of only 2.8 Å. However, no major transition in the receptor-binding domain (RBD) was observed, and the RBD remained in the “Up” position. By measuring the distance of Tyr505His (Protomer B-RBD) with two randomly chosen reference contact residues, Gln134 and His146 in Protomer A, it was found that in the native model, the distance from Tyr505 to Gln134 was 24.35 Å, and to His146, it was 34.78 Å (Figure 3A). In the mutant model, the distance from His505 to Gln134 was 39.87 Å, and to His146, it was 47.26 Å, making the binding pocket more accessible to the hACE2 receptor (Figure 3B). The RMSD plot for native and wild-type conformations showed a relatively higher RMSD, but the mutant model attained stability after 80 ns of simulation with an RMSD < 1 Å (Figure 3C). Fluctuations in the RBD region 438-510 and another peak around the 1038-1140 amino acid residues were observed with RMSF > 1 nm (Figure 3D). The stability of these mutations was also checked through DynaMut2, revealing that His505, Asn950, and Lys969 caused a destabilizing effect with prediction stability scores of −1.22, −0.24, and −0.29, respectively (Table 2). Moreover, these destabilizing mutations also cause intermolecular steric clashes such as Tyr505His causes steric clashes with Val503 and Gly496, Asp950Asn with Leu948, Lys947 and Gly496, Asn969Lys with Ile973 and Ser735 amino acid residues (Figure 4). These results predict that mutations in the wild-type protein leads to changes from the native conformation without clear domain transitions. Overall, the analysis suggests that the spike RBD-based Tyr505His mutation, along with the destabilizing effects of Asn950 and Lys969 mutations in the S2 subunit, may provide better accessibility to hACE2 binding. However, further exploration is needed to test this hypothesis and understand the interaction between the mutant model and the ACE2 receptor/neutralizing antibodies.

**Table 2.** Analysis of S protein stability changes upon mutations.

| AA Native | Position | AA Mutant | Prediction Score (kcal/mol) | Outcome |
| --- | --- | --- | --- | --- |
| **Wild type 7FG7** |  |  |  |  |
| Tyr | 505 | His | -1.22 | **Destabilizing** |
| Asn | 764 | Lys | 0.11 | Stabilizing |
| Asp | 950 | Asn | -0.24 | **Destabilizing** |
| Asn | 969 | Lys | -0.29 | **Destabilizing** |
| **Closed Conformation (6VXX)** |  |  |  |  |
| Tyr | 505 | His | 0.46 | Stabilizing |
| Asn | 764 | Lys | 0.26 | Stabilizing |
| Asp | 950 | Asn | -0.37 | **Destabilizing** |
| Asn | 969 | Lys | -0.4 | **Destabilizing** |
| **Open Conformation (6VYB)** |  |  |  |  |
| Tyr | 505 | His | -1.41 | **Destabilizing** |
| Asn | 764 | Lys | 0.13 | Stabilizing |
| Asp | 950 | Asn | -0.59 | **Destabilizing** |
| Asn | 969 | Lys | -0.49 | **Destabilizing** |

**Figure 3.**
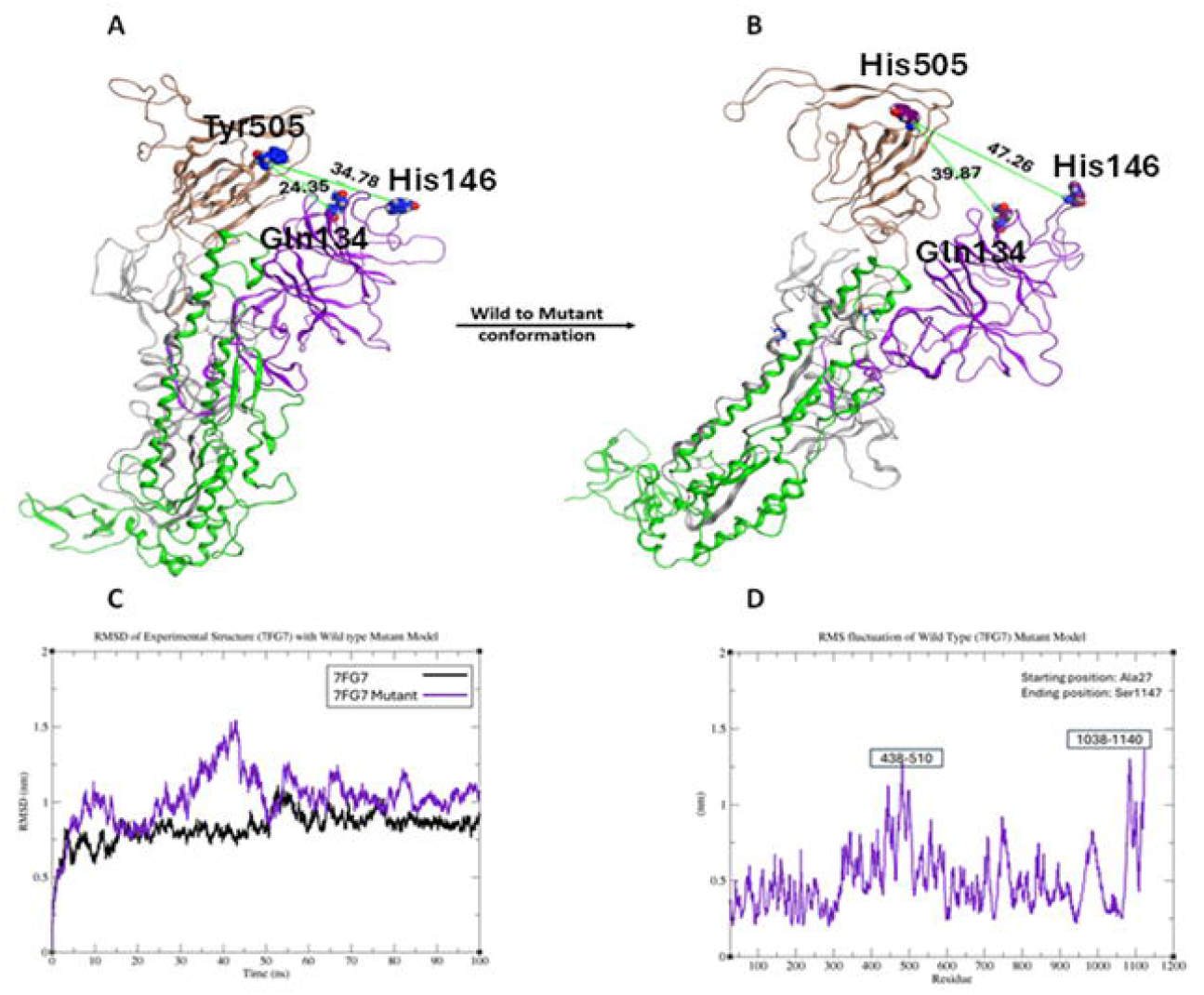
Structural dynamics of native and mutant Wild Type conformation (7FG7) extracted after 100 ns of MD simulation. (A) distance measured between Tyr505 with the reference Gln134 and His146 residues. (B) distance computation of the mutant model with Tyr505 replaced with His505. (C) RMSD plot comparison between native and mutant model of wild type protein. (D) RMSF plot for the wild-type mutant protein

**Figure 4.**
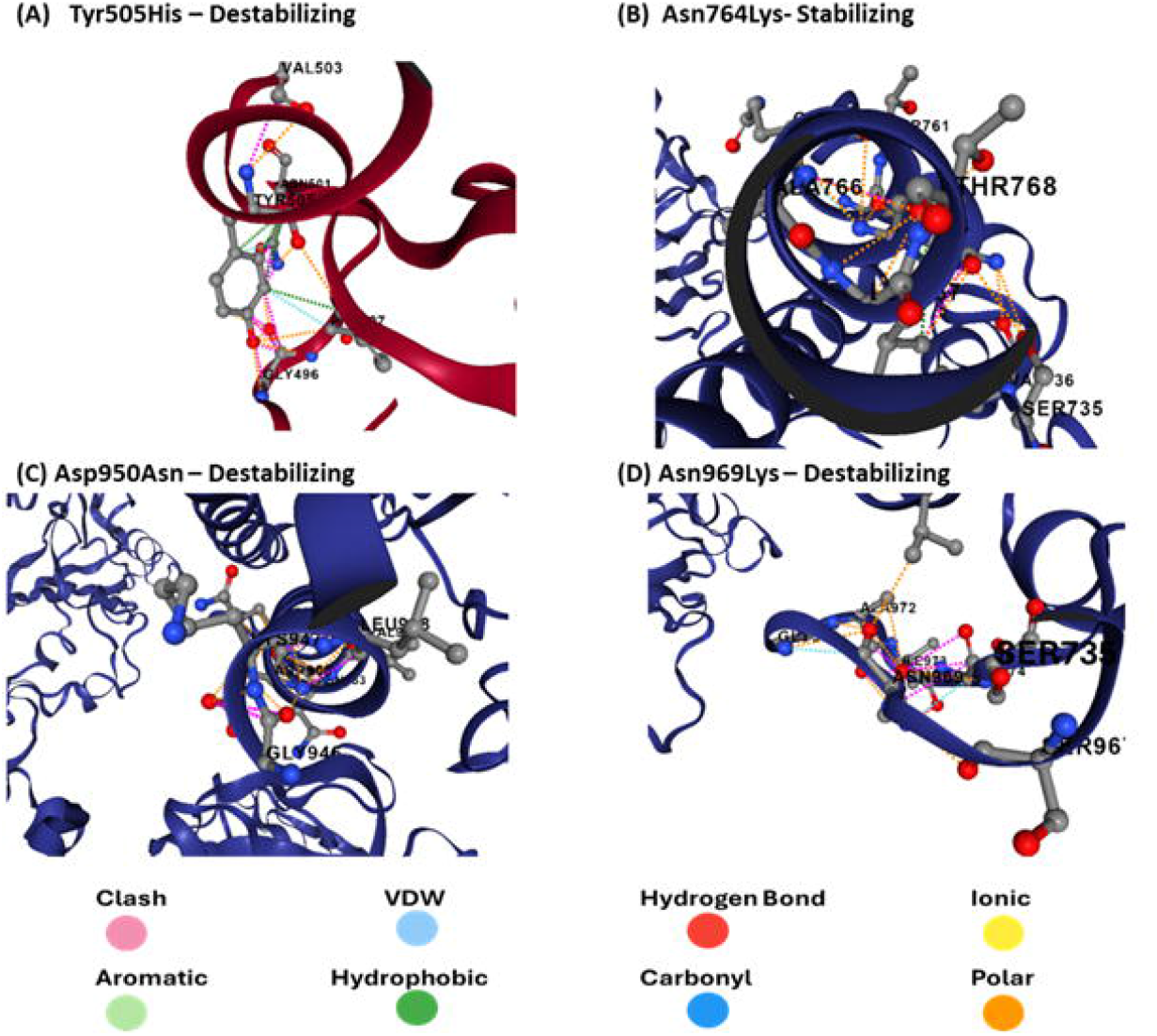
Interaction profiling for Wild Type (7FG7) S protein stability changes upon mutations. (A) destabilizing effect caused by steric clashes (pink dashed bond) upon His505 mutation. (B) Stability interaction profile with no steric clash upon Lys764 mutation. (C) destabilizing effect caused by steric clashes (pink dashed bond) upon Asn950 mutation. (D) destabilizing effect caused by steric clashes (pink dashed bond) upon Lys969 mutation.

In the closed state of the SARS-CoV-2 Omicron spike, mutations introduced in the receptor-binding domain (RBD) region and the S2 subunit induce a structural transition in the α-helix of the S2 subunit protomer C, as shown in Figure 5A. The comparison of the native and mutant model extracted from last frame of MD simulation revealed that these mutations did not cause the RBD to move from the down position to the up position and there was no clear transition in the closed to open state conformation. However, a positional shift in protomer C, which comprises the S2 subunit, was observed. In the native structure, protomer C is in close contact with protomer A, but mutations in the S2 subunit cause it to move towards protomer B (Figure 5A).

**Figure 5.**
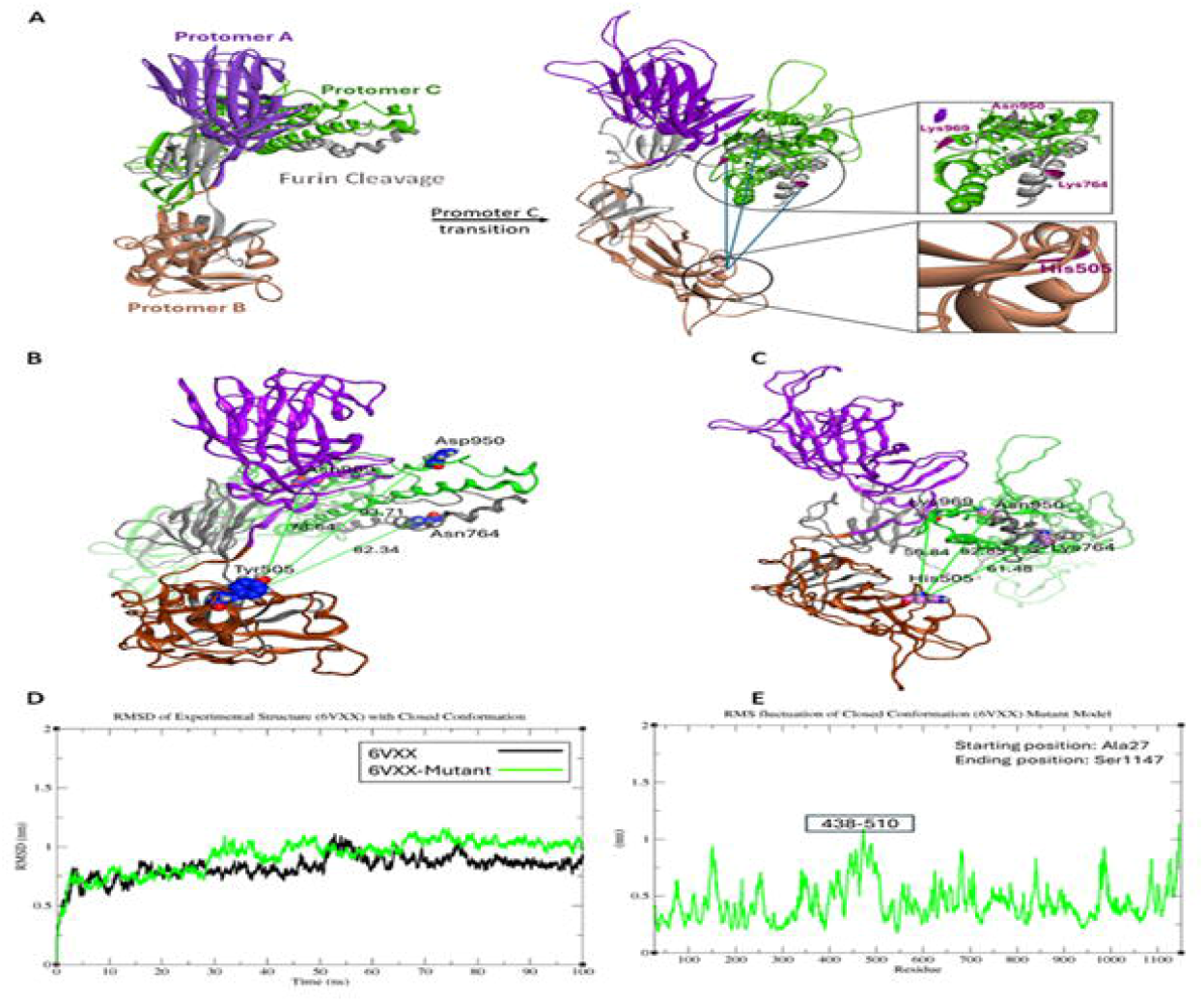
The structural dynamics of native and mutant closed conformation (6VXX) extracted after 100 ns of MD simulation (A) transitional shift of protomer C towards A (B) distance computation of the native model (C) distance computation of the mutant model (D) RMSD plot comparison between native and mutant model (E) RMSF plot for the mutant model

To further validate the movement of protomers, distances were computed using Tyr505His in protomer B (RBD domain) as a reference, alongside other substitutions in the S1/S2 cleavage site (Asn764Lys) and S2 subunit mutations (Asp950Asn, Asn969Lys) in the spike protein. In the native model, the distances between Tyr505 and Asn764, Asp950, and Asn969 were 82.34, 93.71, and 78.64, respectively calculated using distance computation feature from MOE (Figure 5B). Comparatively, in the mutant model, the distances between His505 and Lys764, Asn950, and Lys969 were 61.84, 82.85, and 56.84, respectively (Figure 5C). The reduced distances between the reference His505 in protomer B and other substitutions in the S1/S2 cleavage site and protomer C confirm the movement of protomer C towards the RBD domain in protomer B (Figure 5A-5C).

To further investigate the effect of these mutations, the Root Mean Square Deviation (RMSD) of the native closed conformation (6VXX) and the mutant model was compared after 100 ns of MD simulation. Initially, the RMSD of the mutant model was relatively higher, but after 70 ns, both the native and mutant models reached equilibrium, depicting the stability of both structures (Figure 5D). No significant fluctuations (>1 nm) were observed, except for a minor fluctuation around the 438-510 amino acid residues region of the RBD (Figure 5E). The spike mutation stability change analysis reveals His505 and Lys764 causes stabilizing effect. However, Asn950, and Lys969 caused destabilizing effect with the prediction stability scores of −0.37 and −0.4, respectively (Table 2). The destabilizing effect is mainly caused as discussed earlier due to the steric clashes and in case of closed conformation of spike protein Asn950 causes steric clashes with Leu948 and Lys969 with Gly971 and Ser975 amino acid residues (Figure 6).

**Figure 6.**
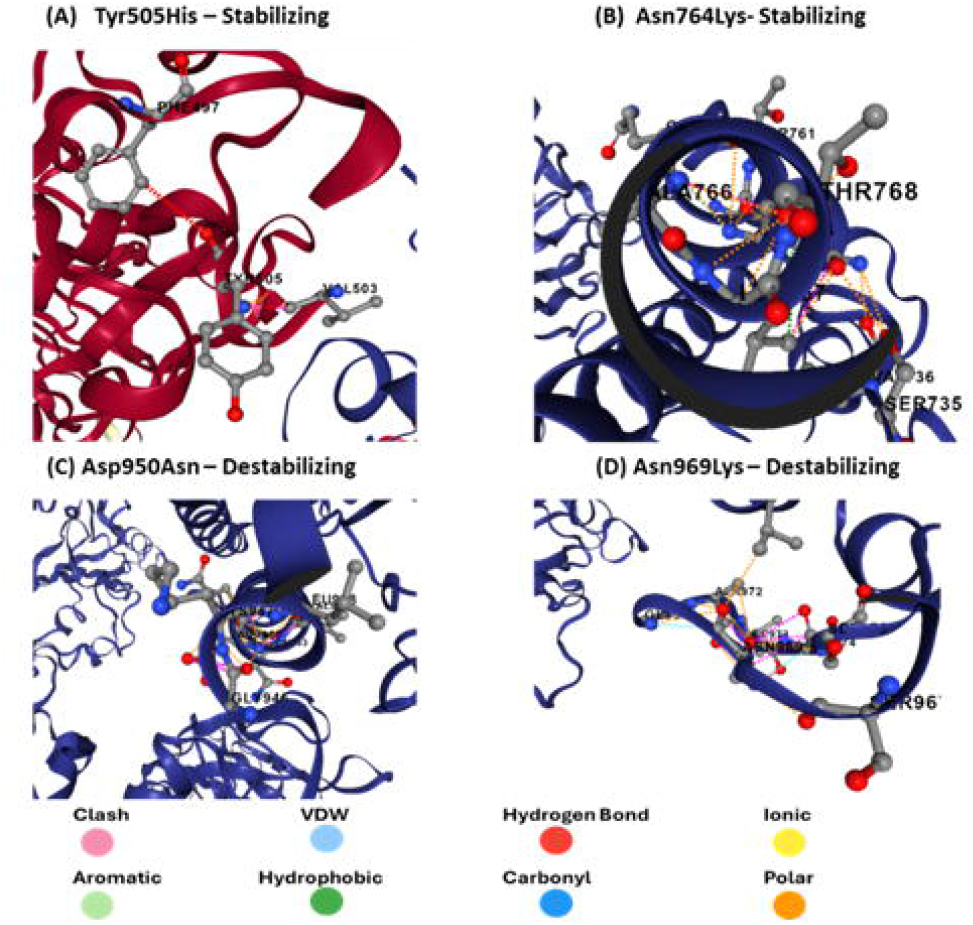
Interaction profiling for Closed Conformation (6VXX) of S protein stability changes upon mutations. (A) Stability interaction profile upon His505 mutation. (8) Stability interaction profile upon Lys764 mutation. (C) the destabilizing effect caused by steric clashes (pink dashed bond) upon Asn950 mutation. (D) destabilizing effect caused by steric clashes (pink dashed bond) upon Lys969 mutation.

In conclusion, the structural transition in the α-helix of the S2 subunit protomer C and the positional shift towards protomer B, induced by mutations in the SARS-CoV-2 Omicron spike, are validated by computed distances. The increased stability observed in the RMSD and RMSF analysis further supports these findings that the conformation indicating positional change of the promoter C towards B, remains stable after 70 ns of MD simulation. This enhanced stability and the reduced accessibility of the RBD due to these mutations may contribute to neutralization resistance against antibodies. Further investigation is required to fully understand the implications of these mutations on the structural dynamics and immune evasion of the Omicron variant.

In the open state, the α-helix of the S2 subunit shifts its position between the domains of the S1 subunit, a crucial binding site for the ACE2 receptor attachment observed in the conformation extracted after 100 ns of the MD simulation. The mutation makes the open conformation more accessible, potentially facilitating viral entry into the host cell. The interdomain interaction pattern in the native model (6VYB), involving RBD amino acids Tyr351, Val401, Asn439, Tyr449, Tyr451, Tyr453, Phe456, Thr470, Tyr505, Gln506, and Pro507 (Figure S3A), is stronger compared to the mutant model (Figure S3B). The transition of the α-helix in the S2 subunit may disrupt hydrogen bonds and π-π interactions. These mutations also assist in trimer opening when we compared them with the wild-type open conformation. We have also observed a sign of structural rearrangement in the S2-α-helix subunit in the region of 946-1010 amino acid residues with spike transition in trimer opening (Figure 7A – 7B). Additionally, the distance between Tyr505 and Gln143 in the binding pocket of the native model is 32.50 Å, while in the mutant model, Tyr505 is replaced by His505, increasing the distance to 37.45 Å (Figure 7C–7D). The mutation in the open conformation increases the accessibility of the ACE2 receptor binding site, potentially enhancing viral infectivity. This suggests that the virus may exploit this mutation to improve its entry into host cells. The increased distance in the mutant model’s binding pocket may alter the binding affinity of the virus to the ACE2 receptor, influencing the efficiency of viral entry.

**Figure 7.**
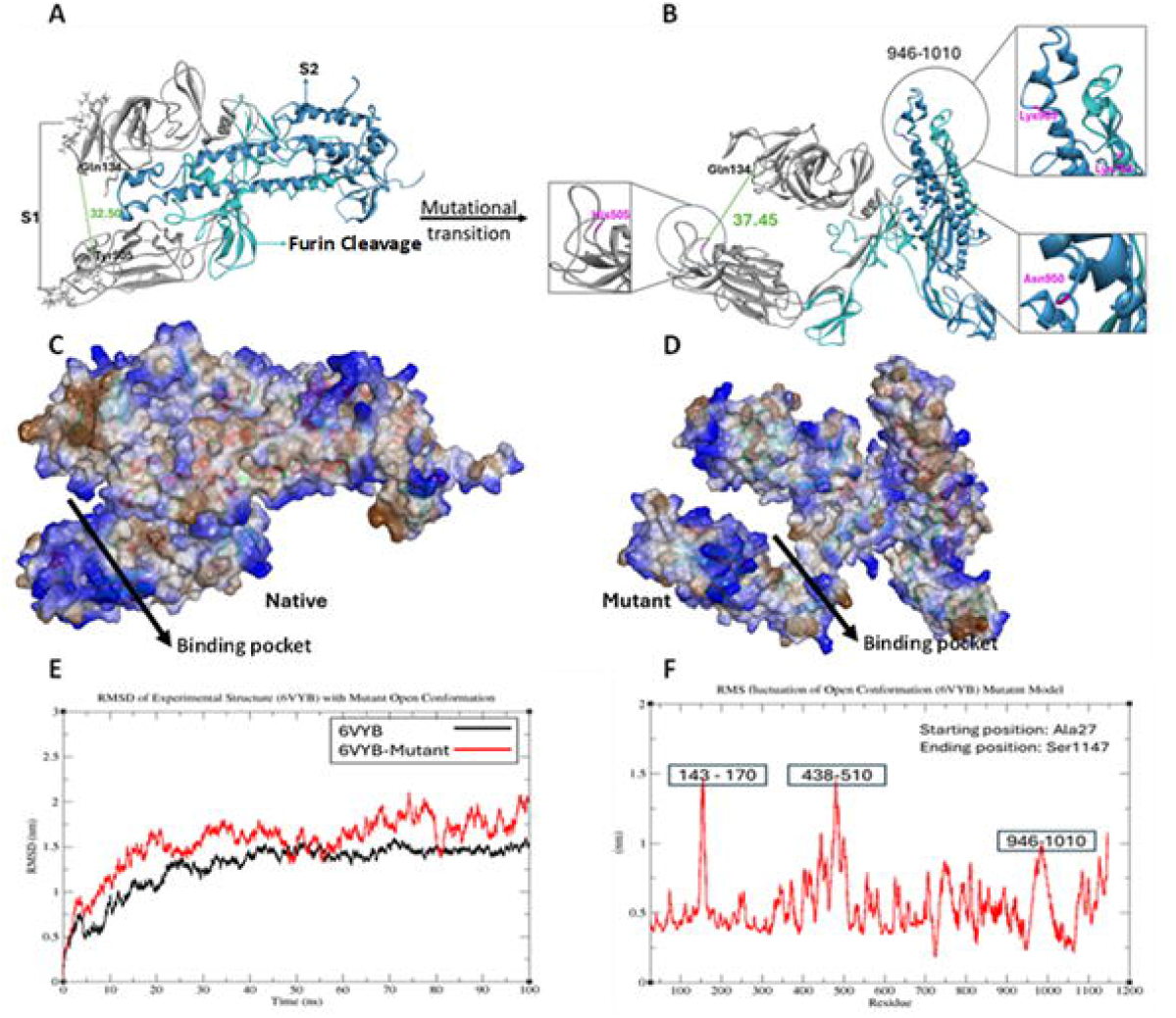
Structural dynamics of native and mutant open conformation (6VYB) extracted from the last frame of 100 ns MD simulation. (A) Native trimeric open conformation. (B) Opening of the trimeric 6VYB receptor in the S2-a-helix subunit by inducing mutations. (C) The binding pocket of the native conformation between the S1 and S2 subunit. (D) Increased surface area of the mutant-6VYB model binding pocket between the S1 and S2 subunit. (E) RMSD profiles of the native and mutant model. (F) RMSF of mutant-6VYB model

The superposition of the native and mutant models of the open conformation shows a significant transition, with RMSD values of 42.5 Å based on the helix and beta sheet Cα atoms. Furthermore, the RMSD trajectories of the mutant model compared with the experimental x-ray crystallographic structure (6VYB) indicate instability in the RMSD plot over 100 ns of MD simulation, whereas the native structure stabilizes after 70 ns of MD simulation (Figure 7E). The higher RMSD values indicate significant conformational changes in the mutant model, which may result in altered functionality. Instability in the RMSD plot further suggests that the mutant structure is less stable than the native, potentially impacting the virus’s ability to maintain its infectious structure over time.

To assess the mobility of the Spike glycoprotein structure, particularly due to the His505 mutation in the RBD domain, the Asn764Lys mutation at the S1/S2 cleavage site, and the Asp950Asn and Asn969Lys mutations in the S2 subunit, we observed two major fluctuations in the S1 subunit around the N-terminal domain (NTD) (143–170 region) and the RBD domain (438–510 region). Continuous fluctuations were also noted in the S2 subunit, with a significant increase around residues 946–1010 (Figure 7F). The increased RMSD and fluctuations in the residues were due to the destabilizing effect of His505, Asn950 and Lys969 with the Predicted Stability Change (ΔΔG Stability) of −1.41 −0.59, −0.49 kcal/mol (Table 2), respectively. The destabilizing effect might be due to the steric clashes of His505 with Gly496 and Val503 (Figure 8A), Asn950 with Gly496 and Leu948 (Figure 8C), Lys969 with Gly971 amino acid residues (Figure 8D). However, Lys764 mutation in the Furin region does not cause any destabilizing effect and no intermolecular steric clashes were observed as shown in Figure 8B. In conclusion, increased mobility and fluctuations in the Receptor Binding Motif (RBM) and S2 subunit of the S protein, particularly due to mutations, suggest that these regions are more dynamic, potentially affecting the virus’s ability to interact with host receptors and immune evasion mechanisms. The steric clashes caused by these mutations promote open conformation of the S protein facilitating in virus entry into the host cell.

**Figure 8.**
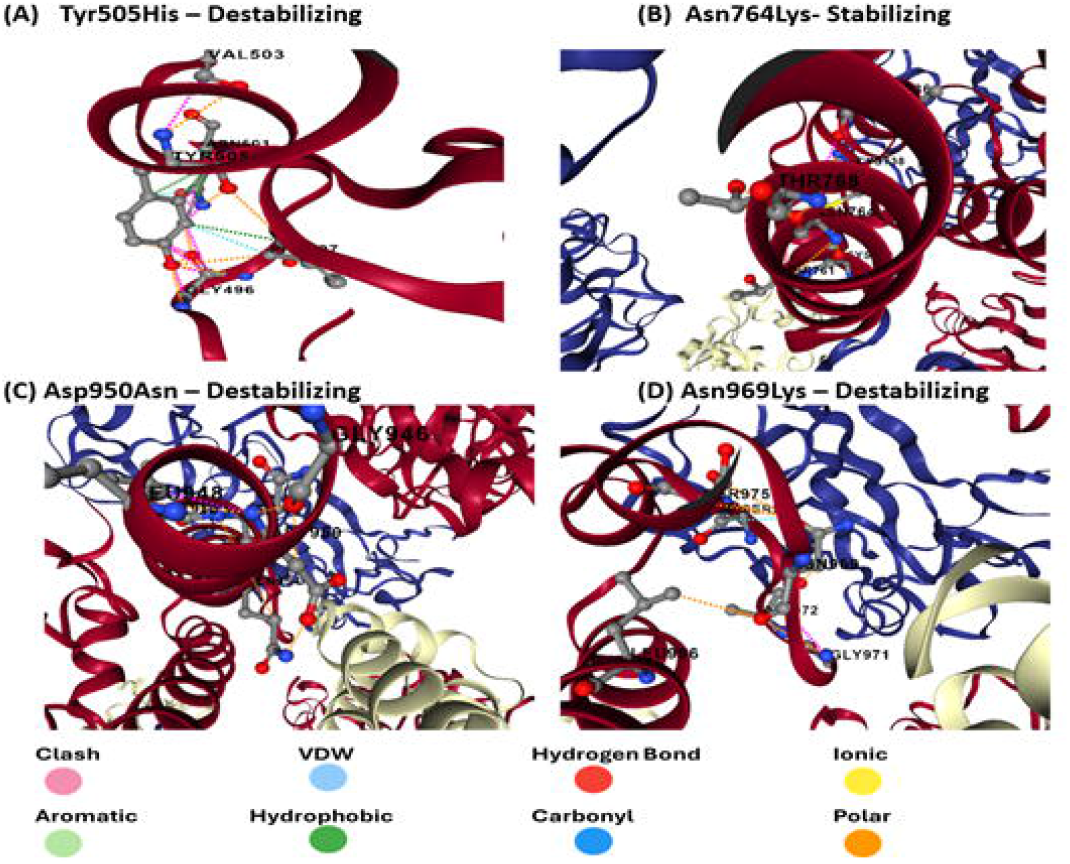
Interaction profiling for Open Conformation (6VYB) of S protein stability changes upon mutations. (A) destabilizing effect caused by steric clashes (pink dashed bond) upon HisSOS mutation. (B) Stability interaction profile with no steric clashed upon Lys764 mutation. (C) destabilizing effect caused by steric clashes (pink dashed bond) upon Asn950 mutation. (D) destabilizing effect caused by steric clashes (pink dashed bond) upon Lys969 mutation.

In addition to studying the wild type, closed, and open conformations independently, we also compared their mutant models based on RMSD graphs generated after 100 ns of MD simulation. The RMSD of the open conformation was found to be higher than that of the wild-type and closed conformations (Figure 9). This aligns with our findings that major structural changes were observed in the open state of the S protein, causing the rearrangement of the trimeric structure. Specifically, the alpha-helix of the S2 subunit moved away from the binding region of the RBD domain in the S1 subunit.

**Figure 9.**
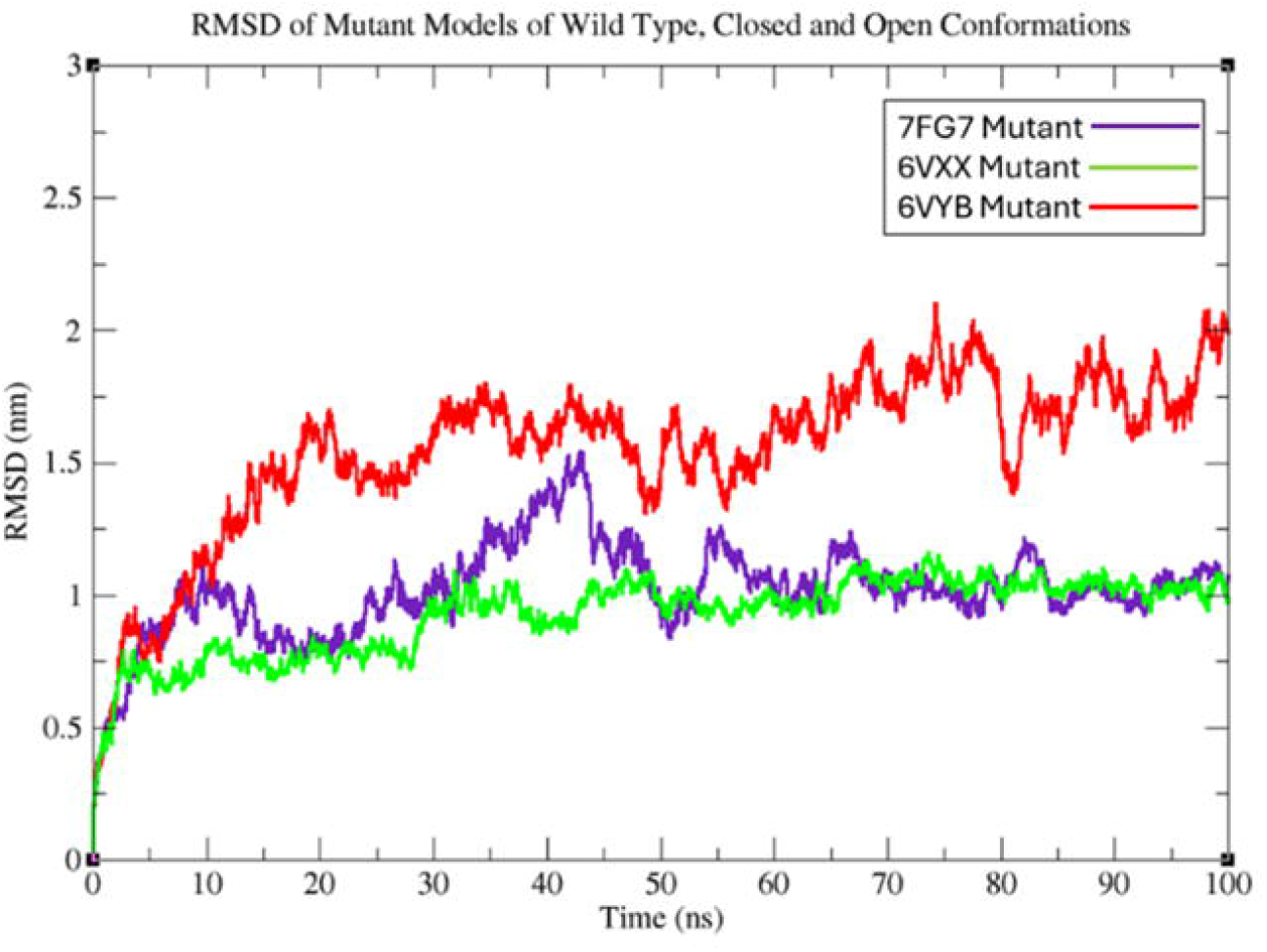
Comparative analysis of Wild-type, close and open conformation mutant models. The indigo trend line represents wild type, green represents closed and red represents open conformation.

## Discussion

The large number of mutations (approx. >30 mutations) observed in the S protein of SARS-CoV-2 omicron variant has raised significant public health concerns [37]. These mutations are present in both S1 and S2 subunits of the S protein [38]. The S1 subunit, which contains the RBM region of the RBD domain, is highly flexible, and mutations in this region cause major structural changes [39]. Mutations in the S2 subunit affect neutralization escape[40], entry route, fusogenicity, and protease requirements[41] Koyama et al. also proposed that the SARS-CoV-2 genome is highly prone to new mutations, which play a significant role in replication and immunological resistance, leading to genetic drift and evasion of immune recognition. Although antigenic drift for SARS-CoV-2 has not been observed, still the virus has the potential to acquire mutations that could facilitate genetic drift and escape [42]. It is the cumulative effect of mutations in both S1 and S2 subunits that contribute to phenotypic changes[41]. In addition, SARS-CoV-2 also causes the evolutionary shifts such as E484K identified in the Beta and Gamma, has been shifted to E484A substitution in omicron, which reflects its dynamic nature[43].

In this study, we investigated the role of S1 and S2 mutations in modulating the structural conformation, opening behavior of RBD in S protein and its accessibility to neutralizing antibodies. Among the 10 deleterious mutations identified, four (His505, Lys764, Asn950 and Lys969) mutations were predominant, and had high VAF (*≥*0.9) as shown in Figure S1. Despite the Omicron variant evolved from Delta, we observed only 1/4 mutations i.e. Asn950 which was also previously reported for Delta[44] was also identified in the omicron variant in our 3 samples. The remaining 03 mutations (His505, Lys764 and Lys969) appear to be unique to Omicron, with no evidence of their occurrence in other VOCs. Interestingly, 3/4 mutations (His505, Asn950 and Lys969) were also found to cause destabilizing effect and contributed to the lower stability of the wild, open and closed state of the S protein as discussed in results.

In the wild-type protein, Tyr505His mutation in the RBM region is being responsible for interaction with ACE2 receptor, as previously discussed by Hossen et al. [45] and Zhang et al. [46]. Zhang also reported that His505 causes to increase the hACE2 binding affinity and ability to escape protective antibodies. It exhibits a functional pattern similar to the well-known E484A substitution in the Omicron RBD. Wang et al. also proposed that both mutations are critical to the RBD region and play comparable roles in influencing receptor recognition, evading RBD-targeting antibodies, and altering entry mechanisms. Lys969 mutation being found in the HR1 region causes destabilizing effect and it has been mainly involved in fusogenicity, compared to Lys764 mutations near the S1/S2 cleavage. It is well understood that the mutations in the HR1 regions (Lys969) are associated with higher fusogenicity[47]. However, it is also important to keep in consideration that fusogenicity caused by the mutations at HR1 region is relatively reduced for omicron (BA.1) and its subvariants versus earlier (B.1.1.7, B.1.351, B.1.6.17.2) variants, which could be one of the reasons for the reduced pathogenicity of Omicron. Moreover, we also found that the Asn950 mutation supports to change the conformation of spike protein, but no functional change was yet to detect in wild-type protein[48].

In the context of closed state, the mutations in RBD (Tyr505His) and S2 subunit (Asn950 and Lys969) increase the binding interface and opening of the RBD binding region, causing the protomer C to move closer to the protomer B (RBD). Furthermore, with regard to open conformation the alpha-helix of protomer C move away from the RBD domain of Protomer B which may aid to reduce interdomain steric hindrance. This movement could create additional space, facilitating the opening of the S trimer. Such structural rearrangements might enhance ACE2 binding and promote the entry of the viral genome into host cells.

Hossen et al. also proposed that the Tyr505His mutation in the wild and closed trimer of S protein can influence the RBD opening [45]. We also demonstrated that the His505Tyr mutation increases the distance between RBD and its binding region, providing a larger binding interface for ACE2 interaction. However, the binding interactions between ACE2 and the mutant models for the omicron variant still needs to be explored. Additionally, we observed that S2 mutations, in combination with S1 (RBD) mutations, help maintain the “down” conformation of RBD. This conformation is likely to enhance antigenic heterogeneity, which may make it more difficult for neutralizing antibodies to recognize the spike protein. Singh et al. reported similar findings, where S2 mutations, in combination with other mutations, increase heterogeneity by retaining the “down” conformation of RBD. This could lead to an increased fraction of the virus population that is not neutralized by antibodies. Consequently, the efficacy of neutralizing antibodies might be reduced, and the retention of S2 mutations could persist, even as the mutational landscape in S1 changes, further complicating immune evasion[15].

To summarize, analyzing RBD-ACE2 interaction in the presence of identified mutations (Lys505His, Asp950Asn, Asn969Lys) in the wildtype, closed and open states is crucial for understanding ACE2 binding patterns. Our study has limitations, focusing on only four mutations and their effects on isolated spike protein models, without examining mutations from other VOCs. Future research should extend this work by investigating the binding affinities of the S protein-hACE2 complex to assess viral attachment and fusion capabilities through molecular modeling and virological approaches.

## Conclusion

In conclusion, the mutations in the RBD (Lys505His) and S2 subunit (Asp950Asn, Asn969Lys) identified from the samples collected in the Karachi region through WGS suggest that these mutations may play a crucial role in the transmission of COVID-19. Moreover, one mutation (Asn764Lys) identified in the Furin region (S1/S2 cleavage site) does not cause any destabilizing effect and no steric hindrances in the intermolecular interaction pattern of S protein. Although these mutations were found in patients with non-severe disease conditions, they still exhibit destabilizing effects in the wild type, closed, and open states of the S protein. Significant conformational changes, particularly in the closed state, where the movement of protomer C towards protomer B (RBD) has been observed. In the open state, displacement of the α-helix in the S2 subunit away from the RBD binding cavity assists in trimer opening. These conformational changes make the receptors more accessible for ACE2 binding. This increased accessibility could facilitate the entry of the viral genome into the host cell, contributing to the infection process. In future studies, the binding interactions between the ACE2 receptor and mutant models containing the Lys505His, Asp950Asn, and Asn969Lys mutations could be analyzed to gain a better understanding of their impact on ACE2 receptor binding.

## Acknowledgement

Field staff, data management unit staff, patients and families for their contribution, CREID network at NIH for funding and technical support.

