## Supplementry tables for "Structural and Conformational Impact of Deleterious Spike Protein Mutations in SARS-CoV-2 Omicron Lineages"

**Table 1:** Statistics of the Mutant Models of Spike (S) Glycoprotein Conformations

| **Model Name** | **Type** | **Dope Score** | **Errat Score** | **Ramachandran Plot** | |
| --- | --- | --- | --- | --- | --- |
|  |  |  |  | **Favorable Region Residues (%)** | **Disallowed**  **Region Residues (%)** |
| 7FG7 Mutant | Wild Type | -123259.0 | 55.5 | 98.5 | 1.5 |
| 6VXX Mutant | Closed Conformation | -116354.2 | 61.47 | 99.3 | 0.7 |
| 6VYB Mutant | Open Conformation | -121739.3 | 65.59 | 99.8 | 0.2 |

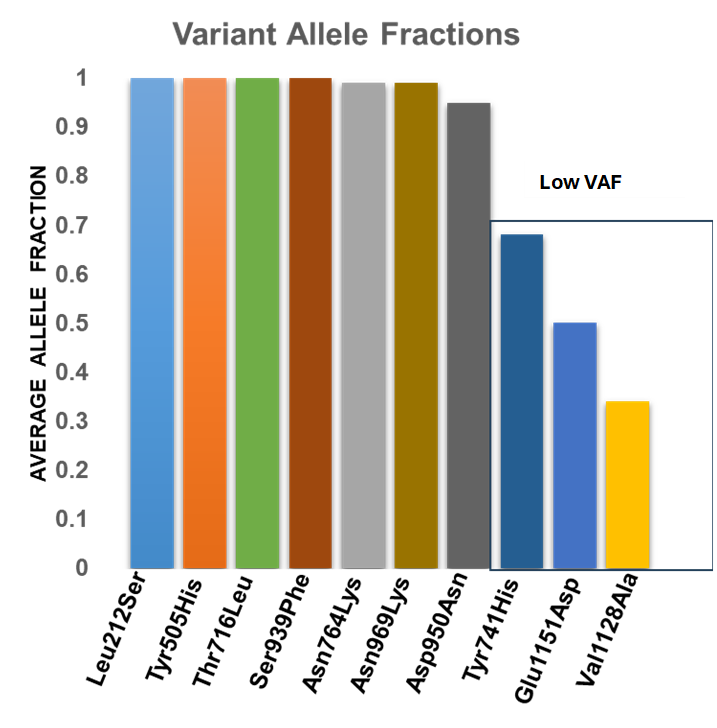

**Figure S1:** Variant allele frequency of the identified deleterious mutations.

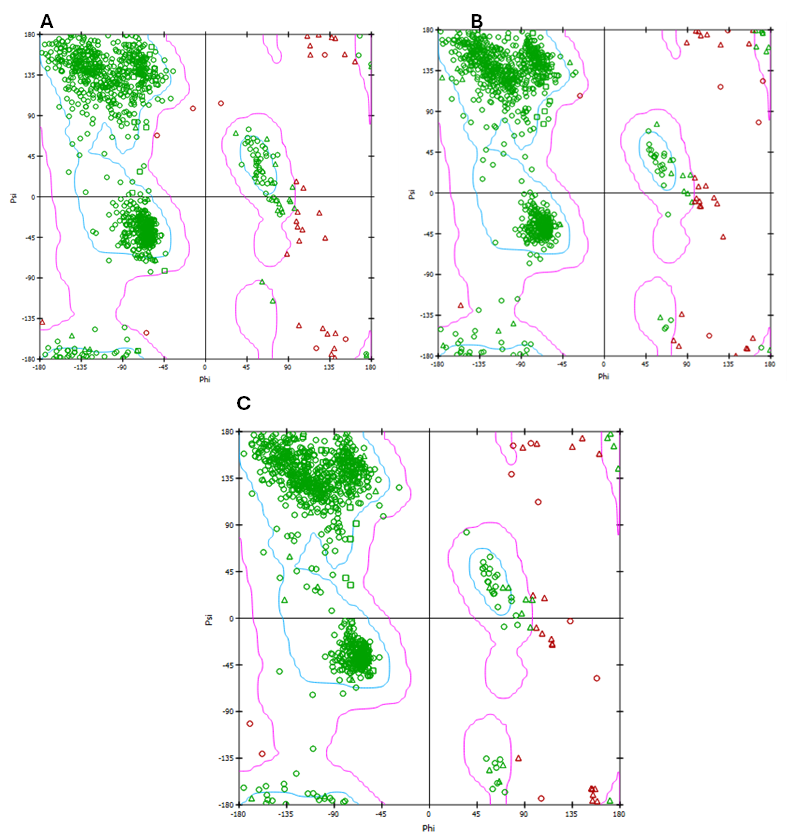

**Figure S2: Ramachandran plot for** (A) Wild type (7FG7) Mutant (B) Closed Conformation (6VXX) mutant model (C) Open Conformation (6VYB) mutant model. Green and pink color represents

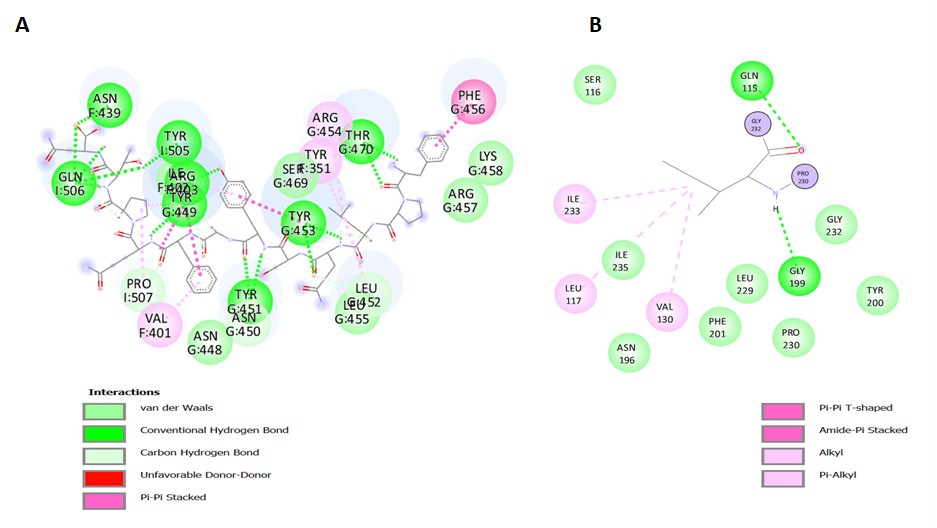

**Figure S3. The intermolecular interaction of open conformation (6VYB) of Spike (S) protein residues.** (A) Wild type (7FG7) (B) Mutant model of 7FG7
